# Distinct Regulation of Host Defences by CRISPR-Cas in Typhoidal and Non-Typhoidal *Salmonella* serovars

**DOI:** 10.64898/2026.02.26.708173

**Authors:** Ankita Das, Nandita Sharma, Sushovan Bhattacharyya, Sandhya Amol Marathe, Vidya Devi Negi

## Abstract

CRISPR-Cas systems are best known for their role in adaptive immunity, but emerging evidence suggests broader regulatory functions. Here, we show that the CRISPR-Cas system acts as a serovar-specific regulator of stress adaptation in *Salmonella enterica*, exerting opposing effects in host-restricted (S. Typhi) and broad-host-range (*S.* Typhimurium) serovars. In *S.* Typhi, CRISPR-Cas system deletion reduces acid and bile tolerance by impairing envelope integrity and repressing key stress-response regulators (*envZ, cadB, phoPQ, lexA, ruvB, wecD*), while increasing resistance to cationic antimicrobial peptides via *pmr* activation and reduced oxidative damage. Conversely, CRISPR-Cas system loss in *S.* Typhimurium enhances acid survival-partly through *speF* upregulation but increases sensitivity to antimicrobial peptides. Spacer-1 of *S.* Typhi CRISPR-I array as the main regulator of gene expression, and its reintroduction partially restored stress tolerance, supporting spacer-dependent control of physiological pathways. These findings establish the CRISPR-Cas system as a non-canonical, spacer-dependent regulator of stress response networks in *S. enterica,* revealing its contribution to the evolutionary divergence of survival strategies between *S.* Typhi and *S.* Typhimurium.

## Introduction

The genus *Salmonella*, a member of *Enterobacteriaceae*, comprises two species-*S. enterica* and *S. bongori*. *S. enterica* is further divided into 6 subspecies and over 2,600 serovars. Clinically, *Salmonella* strains are categorised into two types: typhoidal *Salmonella* (TS) and non-typhoidal *Salmonella* (NTS). Typhoidal serovars are human-adapted pathogens that cause systemic infections (typhoid fever), whereas NTS serovars have a broad host range and primarily cause self-limiting gastroenteritis in humans (Gal-Mor, 2018).The *S.* Typhi and *S.* Typhimurium, TS and NTS serovars, respectively, share approximately 80% genomic similarity, and only about 6% of the shared genes are completely identical (Tang et al., 2013), reflecting their divergent host adaptation and pathogenic mechanisms. Extensive research has explored several pathways and mechanisms to understand the differences in their pathogenesis and response to host stressors (Coluzzi et al., 2025; Garai et al., 2012; Johnson et al., 2018). For instance, *S*. Typhi upregulates the SPI-1 type III secretion system (T3SS) and associated virulence factors to adapt within the gallbladder, thereby promoting chronic carriage, whereas in *S*. Typhimurium, bile downregulates SPI-1 and motility genes, thereby reducing invasion (Johnson et al., 2018). Both survive acidic environments but differ in their acid tolerance; *S*. Typhimurium shows greater tolerance and a greater magnitude of adaptation (Tiwari et al., 2004).

Comparative genomic studies of *S. enterica* subsp. *enterica* highlights the distinct evolutionary trajectory of their CRISPR-Cas system (Kushwaha et al., 2020; Shariat et al., 2015; Pettengill et al., 2014). These studies demonstrate significant divergence in their CRISPR1-leader sequences and *cas* operon sequence and organisation, segregating the *Salmonella* strains into two distinct clades. The clades comprised mainly the TS and NTS serovars, with corresponding CRISPR-Cas variants: CRISPR1-STY/Cas-STY and CRISPR1-STM/Cas-STM. The CRISPR-STY have fewer spacer numbers and diversity. These genomic distinctions could shape host specificity and diversification of virulence among *S. enterica* lineages (Kushwaha et al., 2023, 2020).

Research on the CRISPR-Cas system in various bacteria highlights its significant role in shaping the flow and expression of the accessory genome, underscoring its vital importance in virulence regulation (Yu et al., 2025; Pastuszka et al., 2025; Kushwaha et al., 2020; van Belkum et al., 2015; Lopez-Sanchez et al., 2012). In *S.* Typhi, *cas7* (STY3068) expression has been observed in human macrophages, suggesting a role in intracellular adaptation (Faucher et al., 2006). Additionally, distinct Cas proteins (Cse2, Cas5, Cas6e, Cas1, Cas2, and Cas3) modulate *ompR* expression, thereby influencing porin regulation, bile salt resistance, and biofilm formation (Medina-Aparicio et al., 2021). In non-typhoidal serovars such as *S.* Enteritidis, Cas3 contributes to the regulation of quorum-sensing genes (Cui et al., 2020). Similarly, in *S.* Typhimurium, the CRISPR-Cas system regulates both surface-attached and pellicle biofilm formation (Sharma et al., 2022) and influences its virulence (Sharma et al., 2024). Collectively, these findings underscore the CRISPR-Cas system’s role in modulating virulence-associated processes in *Salmonella*.

Our study aimed to explore the role of the CRISPR-Cas system in regulating the pathogenesis of Typhimurium and Typhi serovars. Given the unique evolutionary trajectories of their CRISPR-Cas systems (Kushwaha et al., 2020) and their role in regulating virulence (Sharma et al., 2024; Cui et al., 2020), we hypothesise that variations in the CRISPR-Cas components may drive distinct bacterial responses to host-associated stressors. We examined how differences in the CRISPR-Cas system influence serovars’ responses to host-associated stressors, including acidic pH, bile salts, and antimicrobial peptides. The results of the study provide insight into how CRISPR-Cas diversity partially shapes *Salmonella’*s adaptation to the hostile host environment. Accordingly, CRISPR-Cas disruption revealed opposing serovar-specific stress phenotypes: exacerbating acid and bile sensitivity in *S.* Typhi while conferring AMP resistance, and enhancing acid tolerance but heightening AMP susceptibility in *S.* Typhimurium. Moreover, spacer-1 of the *S.* Typhi CRISPR-I array emerged as a key regulatory element, displaying the broadest predicted endogenous gene targets and the strongest phenotypic rescue upon reintroduction.

## Material and Methods

### Bacterial Strains and Growth Conditions

*S*. Typhimurium strain 14028s and the *S.* Typhi strain CT18 were used as the parent strain. The wild-type, CRISPR-Cas knockout, and their corresponding complement strains (Supplementary Table 1) were routinely cultured in Luria-Bertani (LB; HiMedia) broth supplemented with appropriate antibiotics and where indicated, 3% (w/v) ox-bile to mimic bile conditions. For assays assessing susceptibility to antimicrobial peptides, bacterial strains were grown in TN medium (0.5% tryptone, 0.5% NaCl) containing polymyxin B at 0.5 µg/mL.

### Construction of the knockout and complement strains

The CRISPR array and *cas3* knockout strains of *S.* Typhi strain CT18 were constructed using a one-step gene replacement strategy outlined by Datsenko and Wanner, 2000 (Supplementary Figure S1). The STYΔ*crisprI* (CRISPR-I array deletion), STYΔ*crisprII* (CRISPR-II array deletion), and STYΔ*cas3* (cas*3* gene deletion) strains were constructed by replacing the target array/gene with a chloramphenicol resistance cassette (Supplementary Figure S2). The primers used for knockout generation, and confirmation are listed in Supplementary Table 2A.

We did not construct a knockout of the *cas*3 gene in *S.* Typhimurium because the *cas3* gene is part of the *cas* operon, and its replacement with a constitutively expressing chloramphenicol cassette would upregulate other *cas* genes, as observed by Cui et al., 2020.

The PCR-amplified fragments of the *cas*3 gene, *crisprI* and *crisprII* arrays (primers are listed in Supplementary Table 2A) were ligated into the pQE60 plasmid (a generous gift from Dipshikha Chakravortty, Indian Institute of Science, India). The verified recombinant plasmids were then introduced into the respective knockout mutants to generate the respective complement strains-STYΔ*crisprI*+pcrisprI, STYΔ*crisprII* + p*crisprII,* and STYΔ*cas*3 + p*cas3* (Supplementary Figure S3). Additionally, STYΔ*crisprI* was complemented with a CRISPR array containing spacer-1 and spacer-4 in sense (STYΔ*crisprI*+p*spacer1*, STYΔ*crisprI*+p*spacer4*) and anti-sense orientation (STYΔ*crisprI*+p*spacer1^r^*, STYΔ*crisprI*+p*spacer4^r^*), and a scrambled array (STYΔ*crisprI*+pscrambled) (Supplementary Table 2B).

### Percentage cell survival under different stress conditions

Overnight-grown bacterial cultures were diluted 1:100 into fresh media containing specific stress factors as described below. All cultures were incubated at 37°C for the indicated durations.

● **Acid tolerance:** Bacterial cultures were sub-cultured in LB medium adjusted to pH 4.0 or pH 7.0 and incubated for 6 h.
● **Bile resistance:** Cultures were grown in LB medium (control) and LB medium supplemented with 3% (w/v) ox-bile for 6 h.
● **Antimicrobial peptide (AMP) susceptibility:** Cultures were sub-cultured in TN medium with or without polymyxin B (PMB) at a final concentration of 0.5 µg/mL and incubated for 7 h.

Following incubation, bacterial suspensions were serially diluted and plated on LB agar containing appropriate antibiotics to determine colony-forming units (CFU). The percentage survival under each stress condition was calculated using the formula:

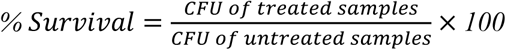

### Cell morphology using field emission scanning electron microscopy

Bacterial cells subjected to bile stress, as mentioned above, were fixed with glutaraldehyde and subsequently dehydrated through a graded ethanol series. The dehydrated samples were air-dried, sputter-coated with gold, and examined using an FEI ApreoS field emission scanning electron microscope (Oxford Instruments, Netherlands).

### Measurement of membrane damage by crystal violet uptake assay

The alteration in membrane permeability was evaluated by crystal violet (CV) uptake assay with some modifications described by Devi et al., 2010. Overnight-grown bacterial strains were subcultured at a 1:100 ratio in LB media with or without 3% ox-bile and grown at 37°C and 150 rpm. After 3 h, bacterial cells were harvested by centrifugation at 5000 rpm for 10 min at 4°C. The cells were washed twice and resuspended in PBS containing 10 μg/ml CV, then incubated at 37°C for 20 min. The suspension was then centrifuged at 13000 g for 15 min, and the OD_590nm_ of the supernatant was recorded. The percentage CV uptake of all the samples was calculated using the following formula:

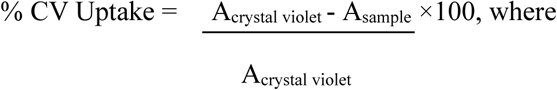

A_sample_-O.D value of the sample and A_crystal violet_-O.D value of the crystal violet solution

### Measurement of reactive oxygen species (ROS) using 2′,7′-dichlorodihydrofluorescein diacetate (DCFDA)

Cells were cultured under the same conditions described previously for bile stress and antimicrobial peptide (AMP) susceptibility assays. After incubation, the cultures were centrifuged at 7000 rpm for 10 minutes, and the resulting pellets were washed twice with sterile phosphate-buffered saline (PBS). The washed cells were then resuspended in sterile PBS and treated with 2 µM DCFDA. The samples were incubated in the dark at 37°C for approximately one hour. Following incubation, fluorescence intensity was measured using a Fluoroskan plate reader (Thermo Scientific) with excitation and emission wavelengths of 485 nm and 535 nm, respectively.

### Measurement of membrane potential

Cells were cultured under the same conditions as described above for AMP susceptibility testing. Following cultivation, the cells were harvested and washed twice with PBS. The washed cells were then resuspended in Milli-Q water and adjusted to OD_600nm_ of 0.6 prior to zeta potential measurement. Zeta potential analysis was performed using a Zetasizer Nano ZS 90 (Malvern Instruments, UK).

### Measurement of hydrophobicity

The absorbance of overnight cultures was measured at OD_600nm_. For the hydrophobicity assay, 3 mL of bacterial suspension was combined with 1 mL of xylene (Himedia), vortexed thoroughly, and incubated at room temperature for 30 min. Following phase separation, the absorbance of the aqueous phase was recorded at 600 nm as described previously (Xu et al., 2009).

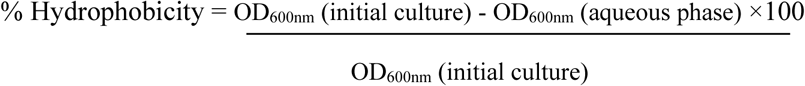

### RNA isolation and quantitative real-time (q-RT) PCR

Bacterial cultures were grown under conditions used to evaluate acid tolerance, AMP susceptibility, and bile resistance, except for bile, where cultures were incubated for 3h. At the end of the specified incubation, the total RNA was isolated using TRIzol reagent (Himedia), and cDNA was synthesised using iScriptTM cDNA synthesis kit (Biorad). Quantitative real-time PCR (qRT-PCR) was performed using iTaq Universal SYBR Green Super-mix (Biorad). Relative transcript levels were determined by the 2^-ΔΔCT^ method, normalising to the reference gene *rpoD*. Primer sequences used in qRT-PCR are listed in Supplementary Table 2A.

### Identification of CRISPR-Spacer Targets

CRISPR-spacer targets were identified through *in-silico* analysis. Spacer sequences from the CRISPR arrays of *S.* Typhi CT18 and *S.* Typhimurium 14028s were aligned against the coding regions of the corresponding parent strains. Reportedly, a 5-bp complementarity between the crRNA and the target DNA is sufficient for Cascade binding to the DNA target (Cooper et al., 2018). Thus, alignments were performed using NCBI BLAST with an expected threshold of 10 and a word size of 5 to obtain the list of potential gene targets.

### Statistical analysis

Statistical analysis was performed using Prism 8 software (GraphPad, California). One-way ANOVA was performed. Error bars indicate standard deviation (SD). Statistical significance is indicated as follows: *, p ≤ 0.05; **, p ≤ 0.01; ***, p ≤ 0.001; ****, p < 0.0001; ns, not significant.

## Results

### CRISPR-Cas system differentially shapes stress responses in Typhoidal and Non-Typhoidal *Salmonella* serovars

To determine whether CRISPR1-STY/Cas-STY and CRISPR1-STM/Cas-STM contribute to divergent pathogenic strategies of *S.* Typhi and *S.* Typhimurium, we compared the susceptibility of the respective wild-type and CRISPR-Cas knockout strains to physicochemical host defences – bile, acidic pH, and AMPs.

In *S.* Typhi, CRISPR-Cas knockout strains showed ∼44.19-71.45% and ∼69.2-84.4% reduction in survival against bile (3% ox-bile) and acid (pH 4) stress, respectively, when compared to wild-type (Figure 1A, Supplementary Figure S4). Conversely, these mutants displayed increased resistance to PMB, with STYΔ*crisprI* and STYΔ*crisprII* strains exhibiting ∼10-fold (≈1000%) higher survival. Although the STYΔ*cas3* strain showed ∼79% lower survival than the wild-type strain, this difference was not statistically significant as per one-way ANOVA with Dunnett’s test (Figure 1A). In contrast, loss of the CRISPR-Cas system in *S.* Typhimurium did not affect bile tolerance but increased survival under acidic stress by ∼42.6-89%. Consistent with previous reports of reduced survival after 1 h PMB exposure (Sharma et al., 2024), treatment for 7 h resulted in a marked decrease (74.23-93.3%) in survival of the *S.* Typhimurium knockout strains (Figure 1B), confirming and extending earlier findings. Complementation of the knockout strains with the corresponding genes restored phenotypes to near wild-type levels, suggesting that the gene deletions are clean with no polar effects.

**Figure 1:**
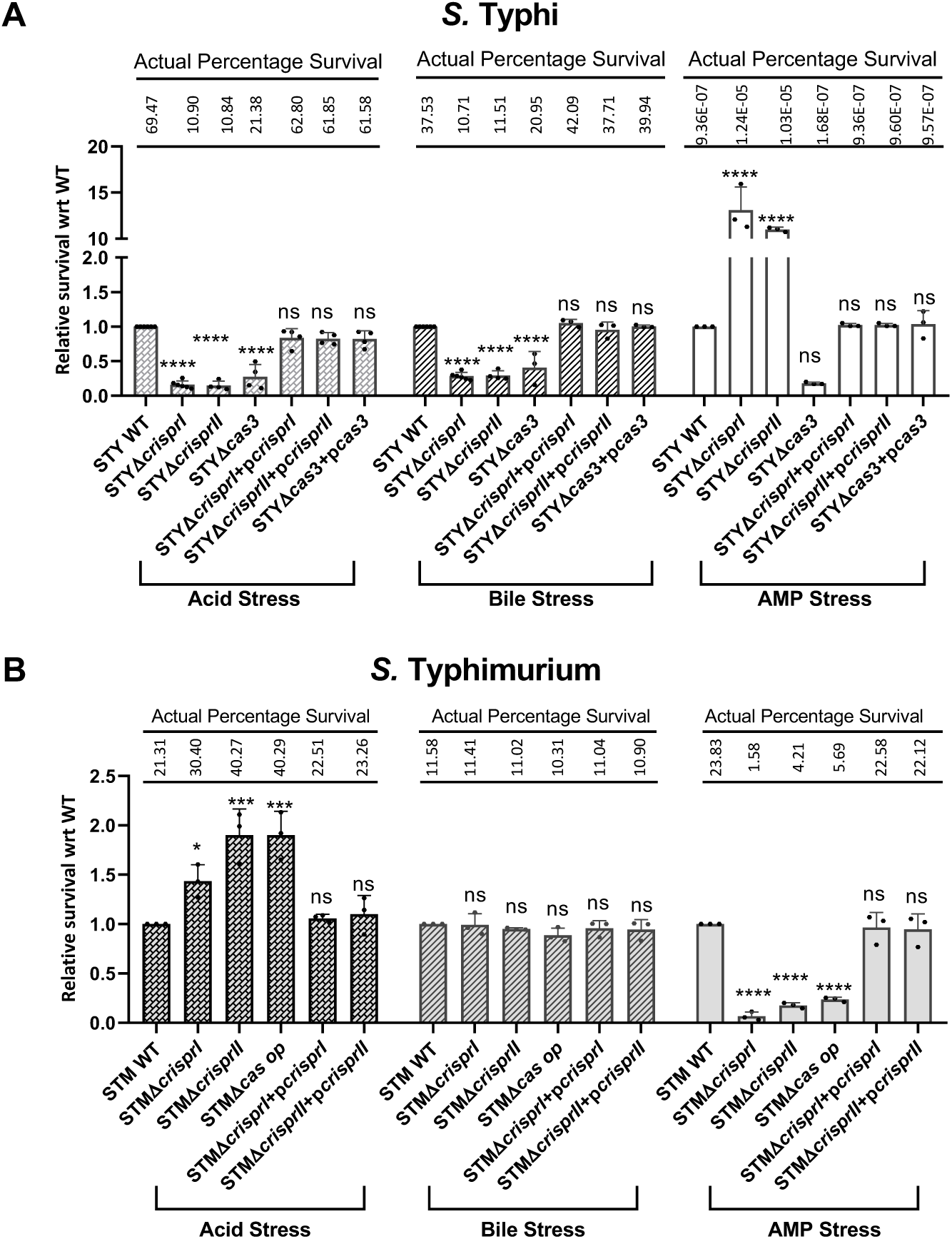
The CRISPR-Cas system contributes differentially to host stress responses, supporting bile and acid resistance in *S.* Typhi, while conferring antimicrobial peptide tolerance in *S.* Typhimurium. The *S.* Typhi strain CT18 wild type (WT), CRISPR (STYΔ*crisprI*, STYΔ*crisprII*), and *cas3* (STYΔ*cas3*) knockout strains, with their complement strains (STYΔ*crisprI*+p*crisprI*, STYΔ*crisprII*+p*crisprII*, and STYΔ*cas3*+p*cas3*) (A), and the *S.* Typhimurium strain 14028s wild type (WT), CRISPR (STMΔ*crisprI*, STMΔ*crisprII*), and *cas* operon (STMΔ*casop*) knockout strains, along with their corresponding complement strains (STMΔ*crisprI*+p*crisprI* and STMΔ*crisprII*+p*crisprII*) (B), were grown under distinct stress conditions. Survival is presented relative to the wild-type strain, while the actual percentage survival for each strain under the specified condition is indicated above the corresponding bars. Data are presented as mean values ± standard deviation. Asterisks indicate statistical significance using one-way ANOVA. *, p ≤ 0.05; **, p ≤ 0.01; ***, p ≤ 0.001; ****, p < 0.0001; ns, not significant. Each experiment was performed with at least three biological replicates (n = 3) and three or more technical replicates (N ≥ 3).

### Oxidative damage does not contribute to reduced survival in *S*. Typhi CRISPR-Cas knockout strains under bile stress, but may contribute to PMB sensitivity

Bile, predominantly composed of bile salts, has been reported to induce oxidative stress in bacterial cells (Walawalkar et al., 2016). To determine whether ROS generation contributed to the reduced survival of *S.* Typhi CRISPR-Cas knockout strains under bile stress, intracellular ROS levels were quantified. No significant differences in ROS levels were detected between the wild-type and CRISPR-Cas knockout strains on bile exposure (Supplementary Figure S5A), suggesting the reduced survival of *S.* Typhi CRISPR-Cas knockout strains under bile stress is not due to oxidative damage. Likewise, no significant difference in bile-induced ROS was observed between the wild-type and CRISPR-Cas knockout strains of *S.* Typhimurium, consistent with their comparable survival patterns (Supplementary Figure S5C).

Interestingly, exposure to PMB revealed serovar-specific differences in ROS production. In *S.* Typhi STYΔ*crisprI* and STYΔ*crisprII* exposed to PMB exhibited ∼50% – 70% lower ROS levels than the wild-type (Supplementary Figure S5B), aligning with their enhanced survival in this condition. Conversely, *S*. Typhimurium, CRISPR-Cas knockout strains exhibited a 13.33% to 26.67% increase in ROS production when exposed to PMB compared to wild-type strains (Supplementary Figure S5D). This elevation in ROS may partially explain the heightened sensitivity of knockout strains to PMB.

### Deletion of CRISPR-Cas exacerbates ox-bile-induced membrane damage in *S.* Typhi, and enhances membrane vulnerability of *S*. Typhimurium to AMPs

Bile exerts antimicrobial activity primarily by disrupting bacterial membranes (Hernández et al., 2012), a mechanism shared with AMPs, which induce membrane damage and lysis (Talapko et al., 2022). Accordingly, bile– and PMB-mediated membrane disruption was quantified using CV uptake assay.

PMB-treated STYΔ*crisprI* and STYΔ*crisprII* showed ∼56-61.3% lower CV uptake than the wild-type (Figure 3A), suggesting reduced membrane disruption, which is consistent with their enhanced survival (Figure 1A). Surprisingly, the CV uptake by PMB-treated STYΔ*cas3* was ∼23.8% less than wild-type (Figure 3A), not matching the survival results. Likewise, PMB-induced CV uptake was similar across all *S*. Typhimurium strains (Figure 3C), despite lower survival of the knockout strains against PMB stress (Fig. 1B).

In *S.* Typhi, CV uptake was measured 3 h post-bile exposure, as extensive membrane damage in *S.* Typhi CRISPR-Cas knockout strains 6 h post-bile exposure precluded reliable quantification. Upon bile exposure, CRISPR-Cas knockout strains exhibited ∼45.8-50.9% higher CV uptake than the wild-type (Supplementary Figure S6A), indicating increased membrane permeability. As expected, *S.* Typhimurium wild-type and knockout strains showed comparable CV uptake 3 h post-bile exposure, consistent with their comparable survival profiles (Supplementary Figure S6B).

Bile-induced structural changes were further examined by scanning electron microscopy. Under LB conditions, all *S.* Typhi strains exhibited intact, smooth outer membranes. Bile exposure caused pronounced morphological damage, including membrane blebbing, surface irregularities, and desiccation (Figure 2A). These were most severe in STYΔ*crisprI* and STYΔ*cas3*, while STYΔ*crisprII* showed intermediate damage. Conversely, bile-treated *S.* Typhimurium wild-type and CRISPR-Cas knockout strains displayed comparable morphologies, like filamentation with limited membrane deformations conforming the ROS-associated stress responses (Figure 2B; de Souza Santos, Salomon and Orth, 2017). Although STMΔ*crisprII* did not undergo filamentation, occasional membrane blebs were observed, but these were not appreciably different from the irregularities seen in the other strains.

**Figure 2:**
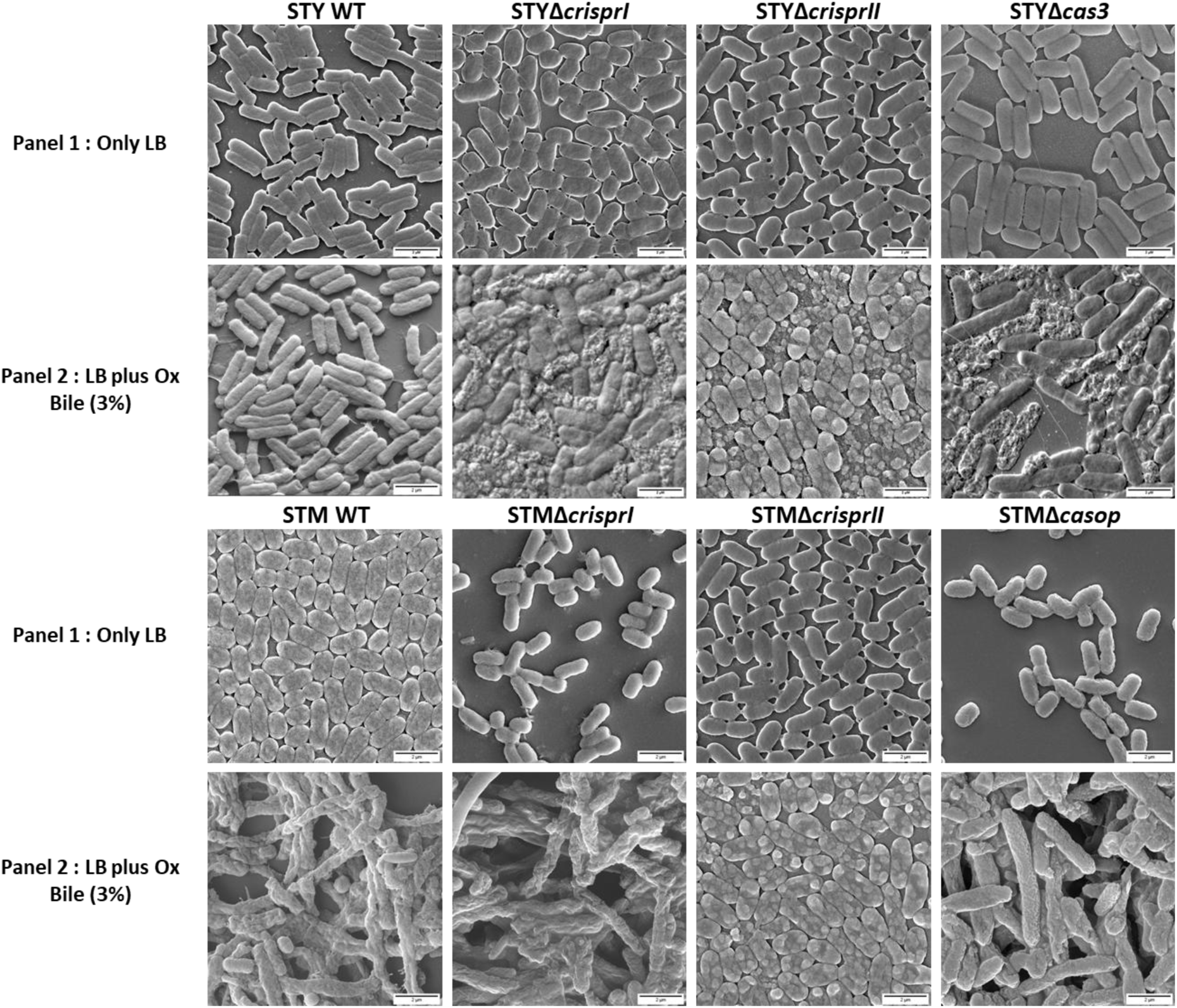
Morphology of bacterial cells under LB and bile stress conditions. Scanning electron microscopy (SEM) analysis was performed to visualise the cell surface morphology of *Salmonella* Typhi wild-type (WT) and knockout (STYΔ*crisprI*, STYΔ*crisprII*, and STYΔ*cas3*), and *Salmonella* Typhimurium wild-type (WT) and knockout (STMΔ*crisprI*, STMΔ*crisprII*, and STMΔ*casop*) strains cultured in LB and LB supplemented with ox-bile (3%) for 6 h. Images were captured at magnifications of 25000×. Scale bar: 2 µm.

**Figure 3:**
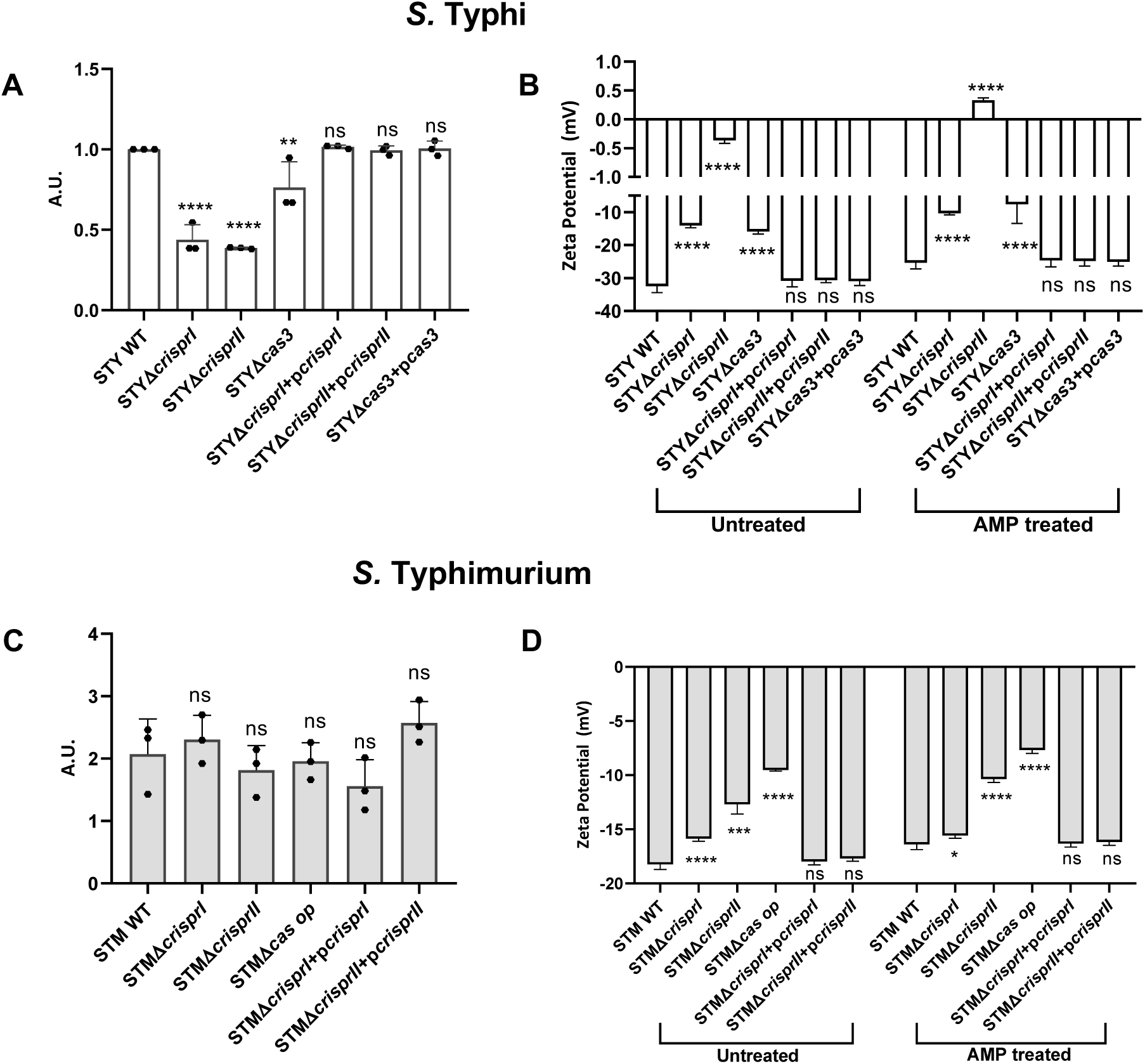
Disruption of CRISPR-Cas components modulates membrane stability and charge regulation in *Salmonella*, influencing resistance to polymyxin B induced stress. *Salmonella* Typhi CT18 wild type (WT), CRISPR knockout mutants (STYΔ*crisprI*, STYΔ*crisprII*) and *cas3* mutant (STYΔ*cas3*), together with their complemented strains (STYΔ*crisprI*+p*crisprI*, STYΔ*crisprII*+p*crisprII*, STYΔ*cas3*+p*cas3*), and *S.* Typhimurium 14028s WT, CRISPR mutants (STMΔ*crisprI*, STMΔ*crisprII*) and *cas* operon mutant (STMΔ*casop*), along with complements (STMΔ*crisprI*+p*crisprI*, STMΔ*crisprII*+p*crisprII*), were analysed for AMP stress-related surface properties. (Figures 3A and 3C) Data are presented as mean ± standard deviation of the ratio of treated to untreated samples for each strain, normalized relative to the wild type. Zeta potential graphs (Figures 3B and 3D) are representative plots. Asterisks indicate statistical significance using one-way ANOVA. *, p ≤ 0.05; **, p ≤ 0.01; ***, p ≤ 0.001; ****, p < 0.0001; ns, not significant. Each experiment was performed with at least three biological replicates (n = 3) and three or more technical replicates (N = 3).

### CRISPR-Cas system regulates surface properties linked to AMP stress adaptation

Cationic modification of lipopolysaccharides (LPS) that increases zeta potential (ZP) of bacteria reduces their susceptibility to AMPs such as PMB (Hwang and Kim, 2024; Olaitan et al., 2014). To investigate whether CRISPR-Cas knockout strains exhibit altered membrane surface charge, we measured ZP in untreated and PMB-treated (0.5 μg/mL) cultures.

PMB treatment moderately increased the ZP of the *S.* Typhi wild-type strain from –32.48 mV to –10.34 mV (Change: 7.132 mV; Figure 3B). The ZP of STYΔ*crisprI* was higher (–14.08 mV) than wild-type, which increased to –10.34mV (Change: 3.74 mV) upon PMB exposure (Figure 3B). The ZP STYΔ*crisprII* strain was the highest (–0.36 mV) and transitioned to a positive value of 0.3324mV (Change: 0.6924 mV) following PMB treatment. The ZP of the STYΔ*cas3* strain increased from –15.88 mV to –7.60 mV following PMB exposure (Change: 8.28 mV). Overall, these trends largely mirrored PMB survival outcomes. However, despite exhibiting similar survival profiles, the STYΔ*crisprI* and STYΔ*crisprII* strains showed markedly different extents of ZP modulation.

PMB treatment caused only minor alterations in the ZP of *S.* Typhimurium strains (Figure 3D). The wild-type strain exhibited a basal ZP of –18.23 mV, which increased slightly following PMB exposure (Change: 1.84 mV). The basal ZPs of all CRISPR-Cas mutants were higher than that of the wild-type, ranging from –15.86 mV to –9.52 mV. Among these, the ZP of the STMΔ*crisprI* strain remained essentially unchanged (ZP of –15.8 mV to –15.5 mV) upon PMB treatment, whereas STMΔ*crisprII* exhibited a small shift (Change: 2.323 mV) and STMΔ*casop* showed a similarly modest change (Change: 1.85 mV). Despite these modest membrane-potential adjustments, all knockout strains displayed reduced survival under PMB stress (Figure 1B).

Membrane-potential changes on PMB exposure were less pronounced in *S.* Typhimurium CRISPR-Cas mutants than those observed in *S.* Typhi CRISPR-Cas knockout strains, highlighting serovar-specific physiological responses. PMB also interacts through its hydrophobic fatty acid tail (Trimble et al., 2016). Cell surface hydrophobicity assessment revealed 98.44% higher hydrophobicity for the STYΔ*cas3* compared to wild-type, whereas all other knockout strains showed reduced hydrophobicity compared to their corresponding wild-type (Supplementary Figure S7).

### Spacer-1 of the CRISPR-I array of *S*. Typhi serves as a primary regulator of the stress response

To explore potential mechanisms underlying CRISPR-Cas-induced alterations in acid, bile, and AMP tolerance, we conducted an *in-silico* analysis to determine whether CRISPR spacers target stress genes and regulate their expression. In *S.* Typhimurium, each spacer had a comparable number of potential gene targets (Supplementary Table 3A), whereas in *S*. Typhi, spacer-1 of the CRISPR1 array had the most targets, some of which are global regulators of the stress response (Supplementary Table 3B). Spacer-6 had the fewest predicted targets, including *ugd*, a key determinant of AMP resistance. Spacer-4 ranked third-lowest in number of predicted target genes (after spacers 6 and 2). Further, STRING network analyses were used to assess indirect links to acid, bile, and AMP stress responses (Supplementary Tables 4B-D). Among the three spacers examined, spacer-4 showed the weakest connectivity to stress-adaptation pathways (Supplementary Table 4A). Thus, spacer-1 of *S*. Typhi was selected for functional complementation of the STYΔ*crisprI*, with spacer-4 and scrambled spacer-1 used as negative controls.

Phenotypic assays showed that complementation of STYΔ*crisprI* with spacer-1 (STYΔ*crisprI*+p*spacer1*) restored acid and bile survival to ∼53.48% and ∼93.1% of the wild-type levels, respectively (Supplementary Figure S7). By contrast, complementation with spacer-4 (STYΔ*crisprI*+p*spacer4*) did not restore survival (Supplementary Figure S8). Notably, complementation with spacers in anti-sense orientation (STYΔ*crisprI*+p*spacer1^r^* and STYΔ*crisprI*+p*spacer4^r^*) had similar results, and the scrambled spacer control (STYΔ*crisprI*+pscrambled) failed to restore survival of the STYΔ*crisprI* to wild-type levels.

The CRISPR II array contains a single spacer with 28 predicted targets, and complementation of STYΔ*crisprII* with this array (STYΔ*crisprII*+p*crisprII*) restored the wild-type phenotype (Figure 1A, and 3-5; Supplementary Figure S8).

### The CRISPR1-STY/Cas-STY and CRISPR1-STM/Cas-STM differentially regulate the stress (acid, bile, and AMP) response genes

Next, expression profiles of potential spacer-targeted genes and other genes involved in the bile/acid/AMP resistance were examined. Expression analysis of acid stress-associated genes revealed significant downregulation of *envZ* (∼1.8-2-fold) and *cadB* (∼2-3-fold) in *S.* Typhi CRISPR-Cas knockout strains relative to wild-type (Figure 4A). Genetic complementation of the knockout strains with full-array or single spacer restored expression of these genes to WT levels, although STYΔ*crisprI*+p*spacer1* showed ∼60% restoration in the gene expression. STYΔ*crisprI*+p*spacer4* behaved similarly to the STYΔ*crisprI*.

**Figure 4:**
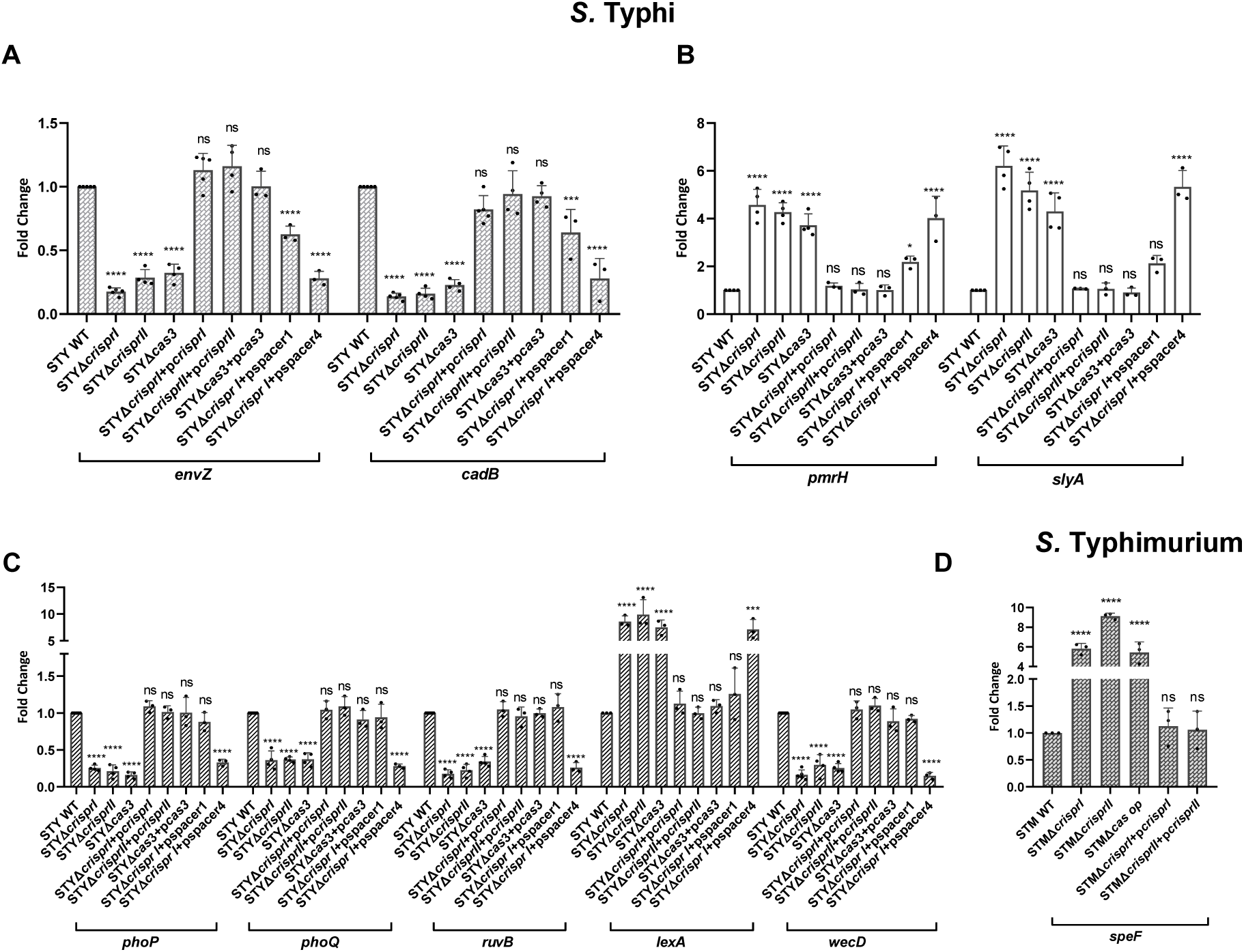
The CRISPR-Cas system exerts differential control over genes mediating acid, cultivated bile, and AMP stress adaptation. *Salmonella* Typhi CT18 (A) wild type (WT), CRISPR knockout mutants (STYΔ*crisprI*, STYΔ*crisprII*) and *cas3* mutant (STYΔ*cas3*), together with their complemented strains (STYΔ*crisprI*+p*crisprI*, STYΔ*crisprII*+p*crisprII*, STYΔ*cas3*+p*cas3*), as well as single spacer complement strains (STYΔ*crisprI*+pspacer1, and STYΔ*crisprI*+pspacer4) were analyzed for expression of (A) acid tolerance genes (*envZ*, *cadB*), (B) AMP resistance genes (*pmrH*, *slyA*), and (C) bile resistance-associated genes (*phoP*, *phoQ*, *ruvB*, *lexA*, *wecD*). (D) In *S.* Typhimurium 14028s WT, CRISPR mutants (STMΔ*crisprI*, STMΔ*crisprII*) and *cas* operon mutant (STMΔ*casop*), along with complements (STMΔ*crisprI*+p*crisprI*, STMΔ*crisprII*+p*crisprII*), expression of *speF* was assessed. Relative gene expression was calculated using the 2⁻^ΔΔ^Ct method, normalized to *rpoD*. Data are presented as mean values ± standard deviation. Asterisks indicate statistical significance using one-way ANOVA. *, p ≤ 0.05; **, p ≤ 0.01; ***, p ≤ 0.001; ****, p < 0.0001; ns, not significant. Each experiment was performed with at least three biological replicates (n = 3) and three or more technical replicates (N = 3).

*In-silico* analysis identified *speF* as a potential CRISPR-Cas target (CRISPR-I_spacer-4, CRISPR-I_spacer-11) in *S.* Typhimurium. Consistent with the enhanced intracellular survival of the CRISPR-Cas knockout strains (Figure 1B), *speF* transcripts were markedly elevated (∼5-9-fold over WT) in these strains (Figure 4B).

The PhoQ regulator, implicated in bile and AMP resistance (Shi et al., 2004; Tsai et al., 2020; van Velkinburgh and Gunn, 1999), was predicted to be a spacer target (*S.* Typhi CRISPR-I_spacer-1; Supplementary Table 3B). At 3 h in nutrient-rich medium, *phoPQ* expression was reduced (∼1.3-2-fold) in *S.* Typhi CRISPR-Cas knockout strains (Figure 4A), while complementation with the corresponding genes restored the expression to wild-type level. Notably, STYΔ*crisprI*+p*spacer1* exhibited expressions similar to wild-type, supporting *phoQ* as a direct spacer-1 target. Bile induces DNA damage and activates the SOS response in *S. enterica* (Prieto et al., 2004). Since *lexA* represses the SOS regulon (Kelley, 2006), and *ruvB*, a DNA repair gene, was predicted as a spacer-5 target, we examined their expression. *lexA* was upregulated ∼7.5-9.9 fold and *ruvB* repressed by ∼1.7-2.7-fold in the knockout strains (Figure 4A). Further, the *wecD* gene, involved in enterobacterial common antigen (ECA) biosynthesis and bile resistance (Bennett and Mitchell, 2025; Ramos-Morales et al., 2003), was reduced by ∼1.3– to 3-fold in knockout strains (Figure 4A). These data indicate global dysregulation of stress response and repair pathways, likely contributing to reduced survival under bile stress.

To explain the altered survival of CRISPR-Cas knockout strains on PMB exposure, lipid A modification genes were analysed. Reportedly, in *S.* Typhimurium CRISPR-Cas knockout strains, increased PMB sensitivity was attributed to downregulation of *pagB*, *pagD*, *pmrD*, *pmrH* (*arnB*), and *pmrE* (*ugd*) (Sharma et al., 2024). As expected, STYΔ*crisprI*, STYΔ*crisprII* showed ∼2-fold upregulation of *pmrH* and *slyA* (Figure 4A), indicating enhanced lipid A remodelling and activation of SlyA-dependent resistance pathways, contributing to their increased survival. However, STYΔ*cas3* showed contradictory results regarding susceptibility to PMB (Figure 1A) and *pmr* gene expression (Figure 4A).

### STYΔ*crisprI* and STYΔ*crisprII* exhibit identical phenotypes owing to repression of the *cas* genes

Although CRISPR-I and CRISPR-II of *S*. Typhi differ in array assembly and spacer-targeting patterns, they displayed nearly identical phenotypes and gene expression profiles. Because CRISPR-Cas components are known to regulate *cas* gene expression (Zakrzewska and Burmistrz, 2023), we analysed *cas* transcription. In both STYΔ*crisprI* and STYΔ*crisprII*, *cas7* and *cas5* were strongly repressed (∼500– and ∼1000-fold, respectively), whereas repression in STYΔ*cas3* was modest (∼3– and ∼16-fold; Figure 5). Likewise, *cas6* expression was markedly reduced (∼50-fold) in STYΔ*crisprI* and STYΔ*crisprII*, while *cse2* showed only moderate repression (∼6-fold); neither gene was significantly affected in STYΔ*cas3*. No significant changes were observed for *cas3* or *cse1* in any knockout strain (Figure 5).

**Figure 5:**
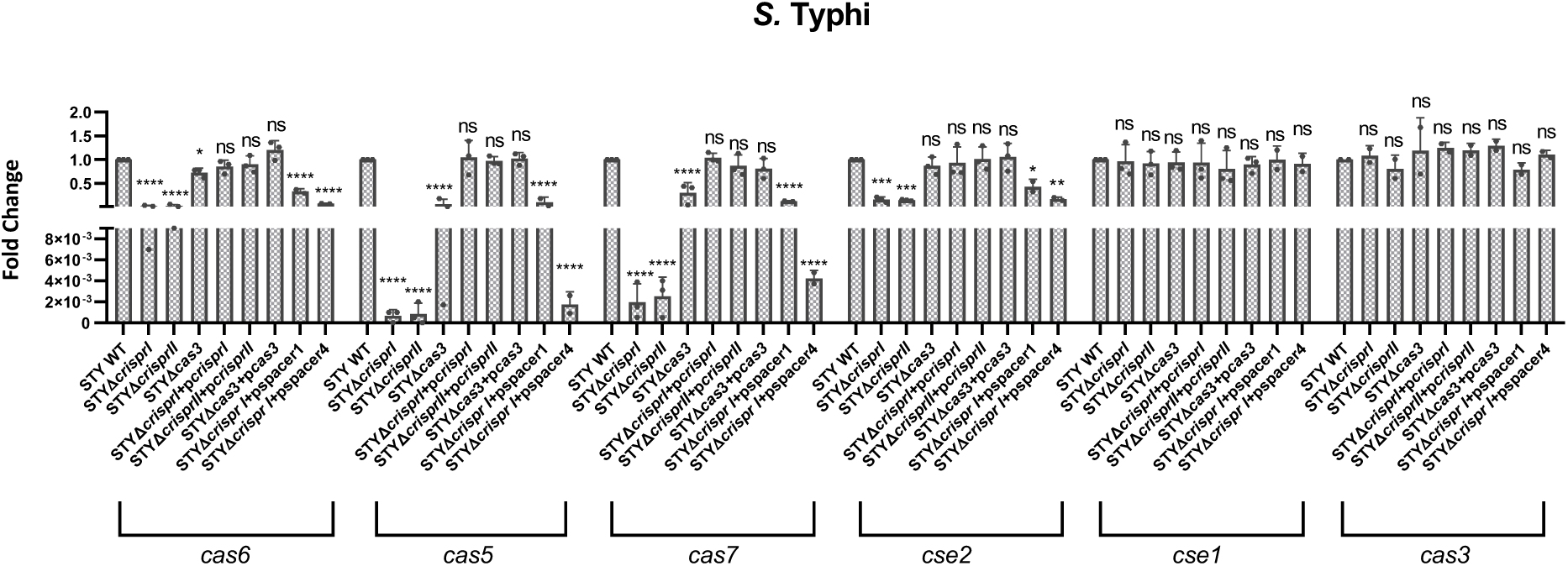
Deletion of the CRISPR-Cas system results in strong repression of *cas* genes expression in *S.* Typhi. In *S.* Typhi CT18 (A), the wild type (WT), CRISPR (STYΔ*crisprI*, STYΔ*crisprII*), and *cas3* (STYΔ*cas3*) knockout strains, with their complement strains (STYΔ*crisprI*+p*crisprI*, STYΔ*crisprII*+p*crisprII*, and STYΔ*cas3*+p*cas3*), as well as single spacer complement strains (STYΔ*crisprI*+pspacer1, and STYΔ*crisprI*+pspacer4) were analysed for expression of cascade genes-*cas6*, *cas5*, *cas7*, *cse2*, *cse1* and *cas3*. Relative gene expression was calculated using the 2⁻^ΔΔ^Ct method, normalised to *rpoD*. Data are presented as mean values ± standard deviation. Asterisks indicate statistical significance using one-way ANOVA. *, p ≤ 0.05; **, p ≤ 0.01; ***, p ≤ 0.001; ****, p < 0.0001; ns, not significant. Each experiment was performed with at least three biological replicates (n = 3), except for strains STYΔc*risprI*+p*spacer1* and STYΔ*risprI*+p*spacer4*, for which two biological replicates were analysed.

**Figure 6:**
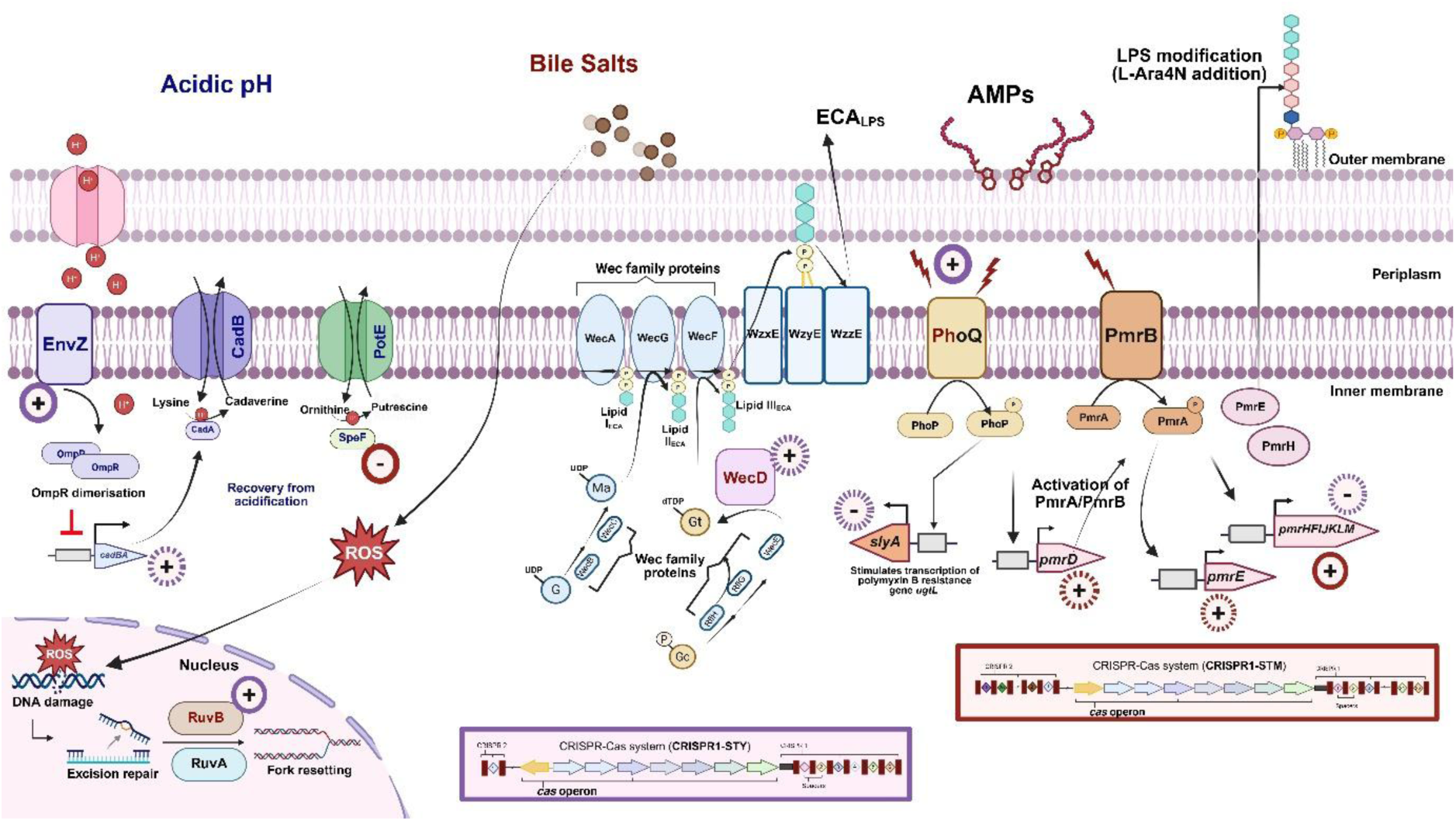
The CRISPR-Cas System Distinctly Regulates *Salmonella* serovars Against Host Defences. Beyond adaptive immunity, the CRISPR-Cas system in *Salmonella* integrates with core regulatory circuits, including two-component systems, LPS remodelling, and DNA repair pathways, to fine-tune bacterial fitness. CRISPR1/Cas-STY promotes survival in hostile host environments by preserving membrane integrity, sustaining genomic repair capacity, and preventing maladaptive overactivation of PhoPQ– and PmrAB-mediated antimicrobial peptide responses. Spacer-1 emerges as a key endogenous regulatory element influencing these processes. In contrast, CRISPR1/Cas-STM exhibits divergent regulatory effects, underscoring serovar-specific evolutionary shaping of physiological control. Collectively, these findings position CRISPR-Cas as a modulator of bacterial physiology and host adaptation rather than solely a defence system. The figure was created using Biorender. Colour coding denotes stress categories: acid (blue), bile (red), and AMP (black). Spacer-mediated regulation is indicated by circled symbols: “+” for positive and “–” for negative regulation. Solid lines represent direct regulation, while dotted lines indicate indirect regulation.

We also examined the expression of *h-ns*, *leuO*, and *crp,* the regulators of the CRISPR-Cas system (Kushwaha et al., 2023). While *h-ns* and *leuO* levels remained unchanged across strains, *crp* was strongly upregulated in STYΔ*crisprI* and STYΔ*crisprII* (∼10-17-fold relative to wild-type) and moderately increased in STYΔ*cas3* (∼2-fold; Supplementary Figure S9). Notably, neither the *cas* genes nor these regulators are predicted spacer targets in *S.* Typhi.

STMΔ*crisprI* and STMΔ*crisprII* also exhibited nearly identical phenotypes and gene expression profiles, likely due to overlapping spacer-targeting patterns. Neither mutant showed significant changes in *cas* gene expression compared to the wild type (Supplementary Figure S10); therefore, regulator expression was not further examined. As expected, STMΔ*casop* showed downregulation of *cas* genes.

## Discussion

*S*. Typhi and *S*. Typhimurium, two serovars of *Salmonella enterica*, employ distinct pathogenic strategies shaped by divergent evolutionary trajectories. *S.* Typhi causes systemic typhoid fever, whereas *S.* Typhimurium induces acute gastroenteritis, differences that extend to their virulence mechanisms, stress responses to bile and acid, and CRISPR-Cas systems. Specifically, *S.* Typhi harbours the CRISPR1/Cas-STY variant, while *S.* Typhimurium encodes CRISPR1/Cas-STM, reflecting their phylogenetic divergence (Kushwaha et al., 2020; Pettengill et al., 2014; Shariat et al., 2015). Adding to these distinctions, our findings indicate that this system contributes to stress adaptation in a serovar-specific manner.

During infection, both serovars encounter acid stress in the stomach and within macrophages *Salmonella*-containing vacuoles. Acid stress activates the EnvZ/OmpR system, promoting OmpR dimerisation and repression of the *cadC/BA* operon, a process essential for cytoplasmic acidification and acid-stress adaptation (Chakraborty et al., 2017, 2015; Rychlik and Barrow, 2005). In *S.* Typhi CRISPR-Cas knockout strains, the pronounced reduction in *envZ* and *cadB* expression suggests disruption of this pathway, likely impairing proton homeostasis and reducing acid tolerance. The gene *envZ* is predicted as a direct target (CRISPR-I_spacer-5), whereas *cadB* is not. Notably, neither *envZ* nor *cadB* is predicted as a direct target of the *S*. Typhi CRISPR-II array. Despite this, complementation of the knockout strains with corresponding CRISPR arrays (STYΔ*crisprI*+p*crisprI* and STYΔ*crisprII*+p*crisprII*) restored the expression of both genes to near wild-type levels. This suggests that CRISPR systems may influence their regulation through mechanisms extending beyond direct spacer-target interactions, like the *cas* gene regulation.

The *spe* gene cluster (*speB, speC, speE,* and *speF*) has been associated with intracellular survival and replication in epithelial cells and macrophages (Schwarz et al., 2022), suggesting an involvement in bacterial acid-stress adaptation. Notably, *speF* (target of CRISPR-I_spacer-4 and CRISPR-I_spacer-11 in *S.* Typhimurium) has been reported to support growth at pH 4.5 (Viala et al., 2011). Its increased expression in the *S.* Typhimurium CRISPR-Cas knockout strains, therefore, raises the possibility that upregulation of *speF* contributes, at least in part, to the improved acid tolerance observed in these mutants.

Upon entry into the intestinal tract, *Salmonella* encounters bile, a complex mixture of bilirubin, cholesterol, and bile salts with potent antimicrobial properties (Giraud et al., 2024; Gonzalez-Escobedo et al., 2011; Hofmann and Hagey, 2008). Bile salts compromise bacterial viability by destabilising membranes, displacing integral membrane proteins, and, following intracellular entry, inducing protein misfolding, denaturation, and DNA damage through their detergent-like activity (Akritidou et al., 2022; Olivar-Casique et al., 2022; Merritt and Donaldson, 2009). Our data suggest CRISPR1/Cas-STY and CRISPR1/Cas-STM have distinct roles in mediating bile resistance. The *S.* Typhimurium CRISPR-Cas mutants showed sensitivity comparable to that of wild-type, while *S.* Typhi CRISPR-Cas mutants displayed pronounced bile sensitivity accompanied by severe outer-membrane damage.

The outer membrane remodelling (Dalebroux and Miller, 2014) and the mechanisms underlying bile resistance in *Salmonella* (Tsai et al., 2020; Prouty et al., 2004) are mediated by the PhoPQ two-component regulatory system. Notably, *phoQ* was identified as a potential target of spacer-1 within the *S.* Typhi CRISPR-I array. Further, the ECA is synthesised by enzymes encoded within the *wec* gene cluster (Bennett and Mitchell, 2025), contributes to membrane integrity and permeability barrier function (Mitchell et al., 2018). Disruption of ECA synthesis, as in *Salmonella* mutants lacking *wecA* or *wecD,* compromises membrane function, resulting in hypersensitivity to bile (Rai and Mitchell, 2020; Ramos-Morales et al., 2003). Consistent with this, reduced expression of *phoP*, *phoQ* and *wecD* in *S.* Typhi CRISPR-Cas knockout strains provides a plausible mechanistic explanation for their compromised membrane integrity and decreased survival under bile stress. Although *wecD* is not a direct target of any spacer within the *S.* Typhi CRISPR arrays, the neighbouring genes *wecE* and *wzxE*, which are co-expressed with *wecD,* are targeted by spacer-1 (STRING analysis; Supplementary Figure S11).

In addition to membrane damage, bile induces genotoxic stress, necessitating efficient homologous recombination and SOS-associated DNA repair pathways (Parmar et al., 2025). We identified *ruvB*, a member of the SOS regulon, as a direct target of CRISPR-I array_spacer-5, linking the CRISPR-Cas system to DNA repair. Reduced *ruvB* expression in CRISPR-Cas mutants suggests impaired recombinational repair and diminished resolution of bile-induced DNA lesions. Consistently, increased *lexA* expression indicates repression of the SOS regulon, providing a mechanistic basis for decreased survival of the mutants under bile stress.

Upon entry into the intestinal lumen, *Salmonella* encounters AMPs that target the negatively charged lipid A moiety of LPS, thereby destabilising the outer membrane (He and Deber, 2024; Hollmann et al., 2018). A common resistance strategy involves lipid A remodelling via the addition of L-Ara4N, which alters surface charge and limits AMP binding (Olaitan et al., 2014). This modification is controlled by the *arnBCADTEF-pmrE* operon (*pmrHFIJKLM-ugd*) and is a hallmark of AMP resistance (Tamayo et al., 2002). Consistent with this model, the improved survival in the STYΔ*crisprI* and STYΔ*crisprII* mutants is associated with increased *pmrH* expression and measurable shifts in cell-surface charge. All CRISPR-Cas knockout strains exhibited elevated expression of *slyA*, a key transcriptional regulator required for full activation of the PhoP-dependent *ugtL* gene, which contributes to AMP resistance (Navarre et al., 2005; Shi et al., 2004). Neither *pmrH* nor *slyA* is a direct spacer target of either CRISPR array. However, *pmrH* is part of the *pmrHFIJKLM-ugd* operon, and *ugd* is a direct target of CRISPR-I_spacer6 and CRISPR-II_spacer1. STRING analysis supports functional connectivity among *pmrH*, *ugd*, and PmrAB pathway components, including *pmrB*, *pmrF*, and *pmrI* (Supplementary Figure S12). Further, the STYΔ*crisprI* and STYΔ*crisprII* show reduced hydrophobicity which could hinder the PMB interaction through its hydrophobic fatty acid tail. Contradictory results were obtained with STYΔ*cas3*, which was susceptible to AMP stress and showed upregulation of *pmrH* and *slyA*. But its increased surface hydrophobicity may promote stronger interactions with the hydrophobic fatty acid tail of PMB, thereby increasing its sensitivity. However, there may be other mechanisms controlling AMP susceptibility in STYΔ*cas3* that require further investigation. *In-silico* analysis shows that *S.* Typhimurium CRISPR spacers target multiple *pmr* genes (*pmrH*, *pmrF*, *pmrK*), linking the CRISPR-Cas system to membrane modification. Consistent with this, disruption of CRISPR-Cas has been reported to reduce *pmr* regulon expression (Sharma et al., 2024), impairing LPS remodelling and adaptation to AMP stress. Notably, the *S.* Typhimurium CRISPR-Cas knockout strains show higher ZP than wild-type, yet they show increased PMB susceptibility with no increase in ZP or CV uptake. Overall, CRISPR-Cas regulates AMP resistance in a serovar-specific manner, promoting *pmr*-dependent protection in *S.* Typhimurium but repressing it in *S.* Typhi.

The *crisprI* and *crisprII* mutants displayed similar phenotypes and gene expression profiles for respective serovars. CRISPR-I and CRISPR-II arrays in *S*. Typhimurium harbour 22 and 26 spacers, respectively, with overlapping gene targets (Supplementary Table 3A), which likely explains the similar phenotypes observed in the knockout strains. However, *S.* Typhi CRISPR arrays have fewer spacers (CRISPR-I:6 and CRISPR-II:1) that have skewed gene targets (Supplementary Table 3B). Further investigation into the basis of this convergence revealed downregulation of genes encoding the Cascade complex in STYΔ*crisprI* and STYΔ*crisprII*, thereby similarly affecting downstream regulatory or stress-response processes. The similar effects were not manifested in STYΔ*cas3,* indicating that the crRNA-Cascade complex binding to target genes primarily influences the expression of the target genes without or with least involvement of Cas3. Further experiments are needed to validate this.

The altered *cas* gene expression profile in STYΔ*crisprI* and STYΔ*crisprII* can be explained through the significant upregulation of *crp*, a repressor of the operon-*cse1*-*cse2*-*cas7*-*cas5*-*cas6e*-*cas1*-*cas2*-CRISPR (Yang et al., 2014). But expression levels of these genes do not follow the expected pattern of decrease with increasing distance from the *cse1* promoter’s transcription start site seen in *E. coli*. However, the *S.* Typhi CRISPR-Cas locus encodes five transcriptional units encoding *cas3*, sense *cse2*, anti-sense *cas2*-*cas1*, anti-sense *cse2*-*cse1*, and *cse1*-*cse2*-*cas7*-*cas5*-*cas6e*-*cas1*-*cas2*-CRISPR that are not regulated by CRP (Medina-Aparicio et al., 2017). Nonetheless, the experimental media used in this study was different from ours – minimal medium versus LB media. As we know that CRP-mediated regulation is highly sensitive to environmental conditions (Zheng et al., 2004), we hypothesise that in LB media *S*. Typhi may modulate CRP-dependent control of the CRISPR-*cas* locus or permit additional regulatory interactions. It may be that CRP-dependent regulation acts differently on these transcriptional units, producing a *cas* gene expression profile that differs from that observed in *E. coli*. Further, *crp* and the *cas* genes are not direct targets of *S*. Typhi CRISPR spacers. Thus, we propose that these genes are indirectly regulated by the crRNA (spacers), possibly by targeting regulatory RNAs that were not included in our gene-target analysis.

Spacer-1 of the *S.* Typhi CRISPR-I array has the widest predicted intra-genomic targets, enriched for genes involved in two-component regulation, stress response, and virulence, supporting a regulatory role. Complementation of STYΔ*crisprI* with spacer-1 partially restored acid tolerance, likely via indirect regulatory effects, while full restoration of bile resistance is consistent with direct targeting of *phoQ*. Additionally, spacer-1 targets *wecE/wzxE* (linked to ECA biosynthesis, Bennett and Mitchell, 2025; Supplementary Figure-S11) and *ruvC* (recombinational repair, Zerbib et al., 1998; Supplementary Figure-S13), suggesting spacer-1’s role in membrane integrity and genome stability under bile stress. Spacer-1also targets *qseB*, a known regulator of the PmrAB system (Olaitan et al., 2014; Supplementary Figure S14), indicating CRISPR-Cas modulates AMP resistance via interconnected regulatory networks. Reintroduction of spacer-4 into STYΔ*crisprI* did not rescue survival or gene expression, consistent with its weak association with stress-adaptation pathways (Supplementary Tables 4B-D). Notably, similar effects were observed when STYΔ*crisprI* was complemented with spacer-1 and spacer-4 in their antisense orientations. This indicates that the CRISPR-Cas-mediated modulatory response occurs at the DNA level, with gene regulation occurring if the spacer binds to either strand. However, this conjecture needs validation. Further, the absence of rescue with the scrambled spacer confirms a sequence-dependent regulation.

In summary, our study reveals that beyond its canonical role as a defence system, the CRISPR-Cas system appears to interface with core regulatory networks, including two-component systems, and pathways governing LPS remodelling and DNA repair to fine-tune *Salmonella* fitness. It exerts a complex, serovar-specific influence on stress adaptation, envelope homeostasis, and antimicrobial resistance. The CRISPR1/Cas-STY supports survival in hostile host environments by preserving membrane integrity, maintaining genomic repair capacity, and constraining maladaptive overactivation of PhoPQ– and PmrAB-dependent responses to antimicrobial peptides. The strong regulatory influence of spacer-1, supported by both *in-silico* predictions and phenotypic rescue experiments, highlights the potential for individual spacers to act as endogenous regulatory elements. In contrast, the divergent responses were observed with CRISPR1/Cas-STM, highlighting that the regulation of physiological responses is shaped by each serovar’s distinct evolutionary trajectory. Together, these insights advance our understanding of CRISPR-Cas systems as modulators of bacterial physiology and host specialisation.

## Acknowledgements

We would like to express our sincere gratitude to Birla Institute of Technology and Science, Pilani, for their invaluable infrastructural support.

## References

1. Akritidou, T., Akkermans, S., Gaspari, S., Azraini, N.D., Smet, C., Van de Wiele, T., Van Impe, J.F.M., 2022. Effect of gastric pH and bile acids on the survival of Listeria monocytogenes and Salmonella Typhimurium during simulated gastrointestinal digestion. Innovative Food Science and Emerging Technologies 82. 10.1016/j.ifset.2022.103161

2. Bennett, H.C., Mitchell, A.M., 2025. Recent advances in understanding of enterobacterial common antigen synthesis and regulation. Open Biol. 15. 10.1098/rsob.250055

3. Chakraborty, S., Mizusaki, H., Kenney, L.J., 2015. A FRET-Based DNA Biosensor Tracks OmpR-Dependent Acidification of Salmonella during Macrophage Infection. PLoS Biol. 13, e1002116. 10.1371/journal.pbio.1002116

4. Chakraborty, S., Winardhi, R.S., Morgan, L.K., Yan, J., Kenney, L.J., 2017. Non-canonical activation of OmpR drives acid and osmotic stress responses in single bacterial cells. Nature Communications 2017 8:1 8, 1587-. 10.1038/s41467-017-02030-0

5. Coluzzi, C., Piscon, B., Dérozier, S., Chiapello, H., Gal-Mor, O., 2025. Comparative genomics of Salmonella enterica serovars Paratyphi A, Typhi and Typhimurium reveals distinct profiles of their pangenome, mobile genetic elements, antimicrobial resistance and defense systems repertoire. Virulence 16. 10.1080/21505594.2025.2504658

6. Cooper, L.A., Stringer, A.M., Wade, J.T., 2018. Determining the Specificity of Cascade Binding, Interference, and Primed Adaptation In Vivo in the Escherichia coli Type I-E CRISPR-Cas System. mBio 9. 10.1128/mBio.02100-17

7. Cui, L., Wang, X., Huang, D., Zhao, Y., Feng, J., Lu, Q., Pu, Q., Wang, Y., Cheng, G., Wu, M., Dai, M., 2020. CRISPR– cas3 of Salmonella Upregulates Bacterial Biofilm Formation and Virulence to Host Cells by Targeting Quorum-Sensing Systems. Pathogens 9. 10.3390/pathogens9010053

8. Dalebroux, Z.D., Miller, S.I., 2014. Salmonellae PhoPQ regulation of the outer membrane to resist innate immunity. Curr. Opin. Microbiol. 17, 106. 10.1016/j.mib.2013.12.005

9. Datsenko, K.A., Wanner, B.L., 2000. One-step inactivation of chromosomal genes in Escherichia coli K-12 using PCR products. Proc. Natl. Acad. Sci. U. S. A. 97, 6640–6645. 10.1073/pnas.120163297

10. de Souza Santos, M., Salomon, D., Orth, K., 2017. T3SS effector VopL inhibits the host ROS response, promoting the intracellular survival of Vibrio parahaemolyticus. PLoS Pathog. 13, e1006438. 10.1371/journal.ppat.1006438

11. Faucher, S.P., Porwollik, S., Dozois, C.M., McClelland, M., Daigle, F., 2006. Transcriptome of Salmonella enterica serovar Typhi within macrophages revealed through the selective capture of transcribed sequences. Proc. Natl. Acad. Sci. U. S. A. 103, 1906–1911. 10.1073/pnas.0509183103

12. Gal-Mor, O., 2018. Persistent Infection and Long-Term Carriage of Typhoidal and Nontyphoidal Salmonellae. Clin. Microbiol. Rev. 32. 10.1128/CMR.00088-18

13. Garai, P., Gnanadhas, D.P., Chakravortty, D., 2012. Salmonella enterica serovars Typhimurium and Typhi as model organisms: revealing paradigm of host-pathogen interactions. Virulence 3, 377–388. 10.4161/viru.21087

14. Giraud, E., Baucheron, S., Foubert, I., Doublet, B., Nishino, K., Cloeckaert, A., 2024. Major primary bile salts repress Salmonella enterica serovar Typhimurium invasiveness partly via the efflux regulatory locus ramRA. Front. Microbiol. 15, 1338261. 10.3389/fmicb.2024.1338261

15. Gonzalez-Escobedo, G., Marshall, J.M., Gunn, J.S., 2011. Chronic and acute infection of the gall bladder by Salmonella Typhi: understanding the carrier state. Nat. Rev. Microbiol. 9, 9–14. 10.1038/nrmicro2490

16. He, S., Deber, C.M., 2024. Interaction of designed cationic antimicrobial peptides with the outer membrane of gram-negative bacteria. Scientific Reports 2024 14:1 14, 1894-. 10.1038/s41598-024-51716-1

17. Hernández, S.B., Cota, I., Ducret, A., Aussel, L., Casadesús, J., 2012. Adaptation and Preadaptation of Salmonella enterica to Bile. PLoS Genet. 8, e1002459. 10.1371/journal.pgen.1002459

18. Hofmann, A.F., Hagey, L.R., 2008. Bile acids: Chemistry, pathochemistry, biology, pathobiology, and therapeutics. Cellular and Molecular Life Sciences 65, 2461–2483. 10.1007/s00018-008-7568-6

19. Hollmann, A., Martinez, M., Maturana, P., Semorile, L.C., Maffia, P.C., 2018. Antimicrobial peptides: Interaction with model and biological membranes and synergism with chemical antibiotics. Front. Chem. 6, 363805. 10.3389/fchem.2018.00204

20. Hwang, D., Kim, H.J., 2024. Mechanisms of Polymyxin Resistance in Acid-Adapted Enteroinvasive Escherichia coli NCCP 13719 Revealed by Transcriptomics. Microorganisms 2024, Vol. 12, 12. 10.3390/microorganisms12122549

21. Johnson, R., Ravenhall, M., Pickard, D., Dougan, G., Byrne, A., Frankel, G., 2018. Comparison of Salmonella enterica serovars Typhi and Typhimurium reveals typhoidal serovar-specific responses to bile. Infect. Immun. 86. 10.1128/IAI.00490-17

22. Kelley, W.L., 2006. Lex marks the spot: The virulent side of SOS and a closer look at the LexA regulon. Mol. Microbiol. 62, 1228–1238. 10.1111/j.1365-2958.2006.05444.x

23. Kushwaha, S.K., Bhavesh, N.L.S., Abdella, B., Lahiri, C., Marathe, S.A., 2020. The phylogenomics of CRISPR-Cas system and revelation of its features in Salmonella. Sci. Rep. 10, 21156. 10.1038/s41598-020-77890-6

24. Kushwaha, S.K., Kumar, A.A., Gupta, H., Marathe, S.A., 2023. The Phylogenetic Study of the CRISPR-Cas System in Enterobacteriaceae. Curr. Microbiol. 80. 10.1007/s00284-023-03298-w

25. Lopez-Sanchez, M.J., Sauvage, E., Da Cunha, V., Clermont, D., Ratsima Hariniaina, E., Gonzalez-Zorn, B., Poyart, C., Rosinski-Chupin, I., Glaser, P., 2012. The highly dynamic CRISPR1 system of Streptococcus agalactiae controls the diversity of its mobilome. Mol. Microbiol. 85, 1057–1071. 10.1111/j.1365-2958.2012.08172.x

26. Medina-Aparicio, L., Rebollar-Flores, J.E., Beltrán-Luviano, A.A., Vázquez, A., Gutiérrez-Ríos, R.M., Olvera, L., Calva, E., Hernández-Lucas, I., 2017. CRISPR-Cas system presents multiple transcriptional units including antisense RNAs that are expressed in minimal medium and upregulated by pH in Salmonella enterica serovar Typhi. Microbiology (United Kingdom) 163, 253–265. 10.1099/mic.0.000414

27. Medina-Aparicio, L., Rodriguez-Gutierrez, S., Rebollar-Flores, J.E., Martínez-Batallar, Á.G., Mendoza-Mejía, B.D., Aguirre-Partida, E.D., Vázquez, A., Encarnación, S., Calva, E., Hernández-Lucas, I., 2021. The CRISPR-Cas System Is Involved in OmpR Genetic Regulation for Outer Membrane Protein Synthesis in Salmonella Typhi. Front. Microbiol. 12, 657404. 10.3389/fmicb.2021.657404

28. Merritt, M.E., Donaldson, J.R., 2009. Effect of bile salts on the DNA and membrane integrity of enteric bacteria. J. Med. Microbiol. 58, 1533–1541. 10.1099/jmm.0.014092-0

29. Mitchell, A.M., Srikumar, T., Silhavy, T.J., 2018. Cyclic enterobacterial common antigen maintains the outer membrane permeability barrier of Escherichia coli in a manner controlled by YhdP. mBio 9. 10.1128/mBio.01321-18

30. Navarre, W.W., Halsey, T.A., Walthers, D., Frye, J., McClelland, M., Potter, J.L., Kenney, L.J., Gunn, J.S., Fang, F.C., Libby, S.J., 2005. Co-regulation of Salmonella enterica genes required for virulence and resistance to antimicrobial peptides by SlyA and PhoP/PhoQ. Mol. Microbiol. 56, 492–508. 10.1111/j.1365-2958.2005.04553.x

31. Olaitan, A.O., Morand, S., Rolain, J.M., 2014. Mechanisms of polymyxin resistance: acquired and intrinsic resistance in bacteria. Front. Microbiol. 5. 10.3389/fmicb.2014.00643

32. Olivar-Casique, I.B., Medina-Aparicio, L., Mayo, S., Gama-Martínez, Y., Rebollar-Flores, J.E., Martínez-Batallar, G., Encarnación, S., Calva, E., Hernández-Lucas, I., 2022. The human bile salt sodium deoxycholate induces metabolic and cell envelope changes in Salmonella Typhi leading to bile resistance. J. Med. Microbiol. 71, 001461. 10.1099/JMM.0.001461

33. Parmar, K., Kumari, Y., Rajmani, R.S., Chakravortty, D., 2025. The resilience of Salmonella to bile stress is impaired due to the reduced efflux pump activity mediated by the antioxidant enzyme YqhD. mSphere 10. 10.1128/msphere.00382-25

34. Pastuszka, A., Mazzuoli, M.V., Crestani, C., Deborde, L., Sismeiro, O., Lemaire, C., Rong, V., Gominet, M., Jacquemet, E., Legendre, R., Lanotte, P., Firon, A., 2025. The virulence regulator CovR boosts CRISPR-Cas9 immunity in Group B Streptococcus. Nature Communications 2025 16:1 16, 5678-. 10.1038/s41467-025-60871-6

35. Pettengill, J.B., Timme, R.E., Barrangou, R., Toro, M., Allard, M.W., Strain, E., Musser, S.M., Brown, E.W., 2014. The evolutionary history and diagnostic utility of the CRISPR-Cas systemwithin Salmonella enterica ssp. enterica. PeerJ 2014, e340. 10.7717/peerj.340

36. Prieto, A.I., Ramos-Morales, F., Casadesús, J., 2004. Bile-Induced DNA Damage in Salmonella enterica. Genetics 168, 1787–1794. 10.1534/genetics.104.031062

37. Prouty, A.M., Brodsky, I.E., Falkow, S., Gunn, J.S., 2004. Bile-salt-mediated induction of antimicrobial and bile resistance in Salmonella typhimurium. Microbiology (N Y). 150, 775–783. 10.1099/mic.0.26769-0

38. Rai, A.K., Mitchell, A.M., 2020. Enterobacterial common antigen: Synthesis and function of an enigmatic molecule. mBio 11, 1–19. 10.1128/mBio.01914-20

39. Ramos-Morales, F., Prieto, A.I., Beuzón, C.R., Holden, D.W., Casadesús, J., 2003. Role for Salmonella enterica Enterobacterial Common Antigen in Bile Resistance and Virulence. J. Bacteriol. 185, 5328–5332. 10.1128/JB.185.17.5328-5332.2003

40. Rychlik, I., Barrow, P.A., 2005. Salmonella stress management and its relevance to behaviour during intestinal colonisation and infection. FEMS Microbiol. Rev. 29, 1021–1040. 10.1016/j.femsre.2005.03.005

41. Schwarz, J., Schumacher, K., Brameyer, S., Jung, K., 2022. Bacterial battle against acidity. FEMS Microbiol. Rev. 46. 10.1093/femsre/fuac037

42. Shariat, N., Timme, R.E., Pettengill, J.B., Barrangou, R., Dudley, E.G., 2015. Characterization and evolution of Salmonella CRISPR-Cas systems. Microbiology (Reading). 161, 374–386. 10.1099/mic.0.000005

43. Sharma, N., Das, A., Nair, A.V., Sethi, P., Negi, V.D., Chakravortty, D., Marathe, S.A., 2024. CRISPR-Cas system positively regulates virulence of Salmonella enterica serovar Typhimurium. Gut Pathogens 2024 16:1 16, 63-. 10.1186/s13099-024-00653-5

44. Sharma, N., Das, A., Raja, P., Marathe, S.A., 2022. The CRISPR-Cas System Differentially Regulates Surface-Attached and Pellicle Biofilm in Salmonella enterica Serovar Typhimurium. Microbiol. Spectr. 10. 10.1128/spectrum.00202-22

45. Shi, Y., Latifi, T., Cromie, M.J., Groisman, E.A., 2004. Transcriptional control of the antimicrobial peptide resistance ugtL gene by the Salmonella Phop and slyA regulatory proteins. Journal of Biological Chemistry 279, 38618–38625. 10.1074/jbc.M406149200

46. Talapko, J., Meštrović, T., Juzbašić, M., Tomas, M., Erić, S., Horvat Aleksijević, L., Bekić, S., Schwarz, D., Matić, S., Neuberg, M., Škrlec, I., 2022. Antimicrobial Peptides—Mechanisms of Action, Antimicrobial Effects and Clinical Applications. Antibiotics 2022, Vol. 11, 11. 10.3390/antibiotics11101417

47. Tamayo, R., Ryan, S.S., McCoy, A.J., Gunn, J.S., 2002. Identification and Genetic Characterization of PmrA-Regulated Genes and Genes Involved in Polymyxin B Resistance in Salmonella enterica Serovar Typhimurium. Infect. Immun. 70, 6770. 10.1128/IAI.70.12.6770-6778.2002

48. Tang, L., Wang, C.X., Zhu, S.L., Li, Y., Deng, X., Johnston, R.N., Liu, G.R., Liu, S.L., 2013. Genetic boundaries to delineate the typhoid agent and other Salmonella serotypes into distinct natural lineages. Genomics 102, 331–337. 10.1016/j.ygeno.2013.07.014

49. Tiwari, R.P., Sachdeva, N., Hoondal, G.S., Grewal, J.S., 2004. Adaptive acid tolerance response in Salmonella enterica serovar Typhimurium and Salmonella enterica serovar Typhi. J. Basic Microbiol. 44, 137–146. 10.1002/jobm.200310333

50. Trimble, M.J., Mlynárčik, P., Kolář, M., Hancock, R.E.W., 2016. Polymyxin: Alternative Mechanisms of Action and Resistance. Cold Spring Harb. Perspect. Med. 6, a025288. 10.1101/cshperspect.a025288

51. Tsai, M.H., Liang, Y.H., Chen, C.L., Chiu, C.H., 2020. Characterization of Salmonella resistance to bile during biofilm formation. Journal of Microbiology, Immunology and Infection 53, 518–524. 10.1016/j.jmii.2019.06.003

52. van Belkum, A., Soriaga, L.B., LaFave, M.C., Akella, S., Veyrieras, J.B., Barbu, E.M., Shortridge, D., Blanc, B., Hannum, G., Zambardi, G., Miller, K., Enright, M.C., Mugnier, N., Brami, D., Schicklin, S., Felderman, M., Schwartz, A.S., Richardson, T.H., Peterson, T.C., Hubby, B., Cady, K.C., 2015. Phylogenetic distribution of CRISPR-Cas systems in antibiotic-resistant pseudomonas aeruginosa. mBio 6. 10.1128/mBio.01796-15

53. van Velkinburgh, J.C., Gunn, J.S., 1999. PhoP-PhoQ-Regulated Loci Are Required for Enhanced Bile Resistance in Salmonella spp. Infect. Immun. 67, 1614–1622. 10.1128/iai.67.4.1614-1622.1999

54. Viala, J.P.M., Méresse, S., Pocachard, B., Guilhon, A.A., Aussel, L., Barras, F., 2011. Sensing and Adaptation to Low pH Mediated by Inducible Amino Acid Decarboxylases in Salmonella. PLoS One 6, e22397. 10.1371/journal.pone.0022397

55. Walawalkar, Y.D., Vaidya, Y., Nayak, V., 2016. Response of Salmonella Typhi to bile-generated oxidative stress: implication of quorum sensing and persister cell populations. Pathog. Dis. 74. 10.1093/femspd/ftw090

56. Xu, H., Jeong, H.S., Lee, H.Y., Ahn, J., 2009. Assessment of cell surface properties and adhesion potential of selected probiotic strains. Lett. Appl. Microbiol. 49, 434–442. 10.1111/j.1472-765X.2009.02684.x

57. Yang, C.D., Chen, Y.H., Huang, H.Y., Huang, H. Da, Tseng, C.P., 2014. CRP represses the CRISPR/Cas system in Escherichia coli: Evidence that endogenous CRISPR spacers impede phage P1 replication. Mol. Microbiol. 92, 1072–1091. 10.1111/mmi.12614

58. Yu, T., Xie, J., Huang, X., Huang, J., Bao, G., Yuan, W., Gao, C., Liu, C., Hu, J., Yang, W., Li, G., 2025. BaeR and H-NS control CRISPR-Cas-mediated immunity and virulence in Acinetobacter baumannii. mSystems 10. 10.1128/msystems.01067-25

59. Zakrzewska, M., Burmistrz, M., 2023. Mechanisms regulating the CRISPR-Cas systems. Front. Microbiol. 14, 1060337. 10.3389/fmicb.2023.1060337

60. Zerbib, D., Mézard, C., George, H., West, S.C., 1998. Coordinated actions of RuvABC in Holliday junction processing. J. Mol. Biol. 281, 621–630. 10.1006/jmbi.1998.1959

61. Zheng, D., Constantinidou, C., Hobman, J.L., Minchin, S.D., 2004. Identification of the CRP regulon using in vitro and in vivo transcriptional profiling. Nucleic Acids Res. 32, 5874. 10.1093/nar/gkh908

